# Sexual dimorphism in the tardigrade *Paramacrobiotus metropolitanus* transcriptome

**DOI:** 10.1101/2024.04.19.590226

**Authors:** Kenta Sugiura, Yuki Yoshida, Kohei Hayashi, Kazuharu Arakawa, Takekazu Kunieda, Midori Matsumoto

**Affiliations:** Faculty of Science and Technology, Keio University 3-14-1 Hiyoshi, Kohoku, Yokohama, Kanagawa, 223-8522, Japan; Institute of Agrobiological Sciences, National Agriculture and Food Research Organization 1-2 Owashi, Tsukuba, Ibaraki, 305-8634, Japan; Institute for Advanced Biosciences, Keio University 403-1 Nihonkoku, Daihojiji, Tsuruoka, Yamagata, 997-0017, Japan; Exploratory Research Center on Life and Living Systems (ExCELLS), National Institutes of Natural Sciences 5-1 Higashiyama, Myodaiji, Okazaki, Aichi, 444-8787, Japan; Department of Biological Science, Graduate School of Science, The University of Tokyo 7-3-1 Hongo, Bunkyo, Tokyo, 113-0033, Japan

**Keywords:** sex dimorphism, tardigrade, genome, transcriptome, *DMRT* gene family, *Paramacrobiotus metropolitanus*

## Abstract

**Background:** In gonochoristic animals, the sex determination pathway induces different morphological and behavioral features that can be observed between sexes, a condition known as sexual dimorphism. While many components of this sex differentiation cascade shows high levels of diversity, factors such as the Doublesex-Mab-3-related transcription factor (DMRT) are highly conserved throughout animals. Species of the phylum Tardigrada exhibits remarkable diversity in morphology and behavior between sexes, suggesting a pathway regulating such dimorphism. Despite the wealth of genomic and zoological knowledge accumulated in recent studies, the sexual differences in tardigrades genomes have not been identified. In this study, we focused on the gonochoristic species *Paramacrobiotus metropolitanus* and employed omics analyses to unravel the molecular basis of sexual dimorphism.

**Results:** Transcriptome analysis between sex identified numerous differentially expressed genes, of which approximately 2,000 male-biased genes were focused on 29 non-male-specific genomic loci. From these regions, we identified two Macrobiotidae family specific *DMRT* paralogs, which were significantly upregulated in males and lacked sex specific splicing variants. Furthermore, phylogenetic analysis indicated all tardigrade genomes lacks the *doublesex* ortholog, suggesting *doublesex* emerged after the divergence of Tardigrada. In contrast to sex-specific expression, no evidence of genomic difference between the sexes were found. We also identified several anhydrobiosis genes exhibiting sex-biased expression, possibly suggesting a mechanism for protection of sex specific tissues against extreme stress.

**Conclusions:** This study provides a comprehensive analysis for analyzing the genetic differences between sexes in tardigrades. The existence of male-biased, but not male-specific, genomic loci and identification of the family specific male-biased *DMRT* subfamily would provide the foundation for understanding the sex determination cascade. In addition, sex-biased expression of several tardigrade-specific genes which are involved their stress tolerance suggests a potential role in protecting sex-specific tissue and gametes.

## Introduction

Reproductive modes in animals are typically categorized into two major categories: asexual and sexual. Sexually reproducing animals produce sex-specific gametes, and genetic exchange between sexes leads to higher genetic diversity (1, 2). Gonochoristic animals usually show sexual dimorphism not only in gametes but also in somatic tissues, physiology, and behavior within a single species, demonstrating the dynamic differentiation observed in intraspecies.

Although many aspects of the mechanism for inducing sex differences remain to be elucidated, they are usually regulated by sex differences in the genome and gene expression (3). Gonochoristic animals must undergo sex determination through common well-studied mechanisms to develop sex-specific organs. The physiological systems of sex determination vary among species but are generally categorized into two types: determination by sex-linked chromosomes and environmental cues (4, 5). Species possess sex chromosomes that show different karyotypes depending on their sex. In contrast, environmental cues, including temperature, nutritional status, and population density, act as initial cues for sex determination (6). Regardless of the mode of sex determination, several widely conserved genes play crucial roles in sex-specific organ development. The transcription factor family Doublesex and Mab-3 Related Transcription Factor (DMRT) is a key regulator of somatic tissue development in various animals (7). In animals utilizing sex chromosomal sex determination systems, not only *DMRT* orthologs on the sex chromosome, but also on the autosomes are involved in sex determination cascades to regulate the growth of sex-specific tissues (8, 9). In contrast to the chromosomal sex determination system, environmental cues induce the development of sex-specific tissues in normally parthenogenetic individuals through the expression of *DMRT* orthologs (e.g., *Dsx1* in *Daphnia magna*), leading to genetic exchange through mating (10-12).The interplay between highly diverse and conserved components in generating different sexes to overcome environmental and genetic challenges presents a significant challenge for understanding the sex determination cascade.

The phylum Tardigrada, a member of the Ecdysozoa with 1,500 estimated species (13), is divided into three classes: Heterotardigrada, Eutardigrada, and *nomen dubium* Mesotardigrada (14, 15). Tardigrades are renowned for their ability to tolerate extreme environments, and studies have identified tardigrade-specific proteins that mediate tolerance against nearly complete desiccation and anhydrobiosis (16). Asexual (parthenogenesis) and sexual reproduction have been observed within this phylum, with reported instances of both gonochorism and hermaphroditism in sexually reproducing species (17). Sexual dimorphism in morphology and behavior during mating have been observed (18, 19). In contrast, we lack knowledge on the molecular mechanisms that induce sexual dimorphism because most molecular and genomic studies have focused on parthenogenetic species (20).

To this end, we conducted genomic and transcriptomic comparisons between males and females of the model gonochoristic tardigrade *Paramacrobiotus metropolitanus* to identify the molecular factors related to sexual dimorphism. This species, which is rich in ecological information, has a reported 170 Mbp genome, is relatively easy to culture, and show a male-biased sex ratio (Male:Female=7:3), but morphological sexual dimorphism excluding testis/ovary has not been described (19, 21-24). The results in this study lay the foundation for subsequent studies aimed at identifying a master regulator of the sex determination cascade and sex-dependent genetic differences in tardigrades.

## Methods

### Tardigrade culture condition and specimen preparation

The tardigrade *P. metropolitanus* TYO strain was cultured following methods described in the previous report (23). The specimens were sexed using the method described by Sugiura *et al.* (21). The eggs of *P. metropolitanus* were individually placed in an agar-coated dish, and hatched individuals were separated and reared separately to avoid sex contamination. These specimens were then grown until the development of sexual organs that were used for sexing.

### RNA extraction and sequencing

Total RNA was extracted as described by Arakawa *et al.* (25). Two hundred and fifty specimens of each sex were placed in a 1.5 ml tube with minimal water, and 100 µL of TRIzol reagent was added (Thermo Fisher Scientific). Total mRNA was extracted using the Direct-zol RNA kit (Zymo), and the samples were transported to Chemical Dojin for sequencing. The transcriptome sequencing libraries were prepared with poly A selection using the NEBNext Ultra II RNA Library Prep Kit for Illumina (New England Biolabs) and were sequenced using the NovaSeq 6000 instrument (Illumina, 150 bp PE). Four and three replicates were prepared for males and females, respectively.

### External data and annotation

Genome data for *P. metropolitanus* were obtained from our previous study (22). Raw gDNA-Seq reads used to assemble the genome and RNA-Seq data for the hydrated and desiccated samples (2d) were downloaded from SRA with prefetch and fasterq-dump from the sra-toolkit suite v2.10.1 (https://trace.ncbi.nlm.nih.gov/Traces/sra/sra.cgi?view=software, Accession ID: DRR144969, DRR146886). We have added additional annotations to the protein sequences using NCBI Conserved Domain Search (26), DeepLoc2 (27), or InterproScan v5.62-94.0(28). Tardigrade specific anhydrobiosis genes were annotated based on previous studies (22, 29-32). Nucleotide sequences for the coding regions were extracted using gffread v0.12.7 (33). Protein structures were predicted by ColabFold2 v1.5.3 (https://colab.research.google.com/github/sokrypton/ColabFold/blob/main/AlphaFold2_complexes.ipynb) (34) with default settings and visualized ChimeraX v.1.7.0 (35). The chromosome-level genome of *Hypsibius exemplaris* was downloaded from DNAzoo (36) and the positions of the gene predictions from our previous study (31) were converted to the new genome with LiftOff v1.6.3. Genome and gene predictions for *Ramazzottius varieornatus* were obtained from our previous report (31).

### Gene expression analysis

Raw RNA-Seq reads were mapped to the coding sequences and quantified using RSEM v1.3.3 (37). The raw counts were then subjected to statistical testing using DESeq2 v1.38.0, within the run_DE.pl from the Trinity pipeline v2.15.1 (38, 39). Transcripts with FDR values < 0.05 were identified as differentially expressed genes (DEGs). Gene Ontology Enrichment Analysis (GOEA) was performed based on InterProScan GO annotations using GOstats v2.68.0 and GSEABase v1.64.0 (40, 41). Gene ontologies with *p*-values < 0.05 were considered significant. Singleton terms were removed from the final list of enriched terms.

To extract genomic regions enriched in transcripts biased to either sex, we performed enrichment analysis based on the number of DEGs with more than 10x fold change within 200 kbp windows (100 kbp steps). Genomic bins were created with BEDtools v2.31.1 (42) and the number of genes fitting the criteria was calculated using BEDtools intersect. An in-house Rscript was used to perform Fisher’s exact test for each bin, and *p*-values were corrected by BH method. Regions with Q-value < 0.01 were considered enriched.

We also performed transcriptome assembly through Trinity v2.15.1 and StringTie v2.2.1 (39, 43). The RNA-Seq data were mapped to the genome using Hisat2 v2.1.0 (44) and assembled with genome-guided Trinity or StringTie. A non-genome-dependent assembly was also produced with Trinity. The assembled information was passed into PASA v2.5.3 for variant detection and merged with the original gene prediction using EvidenceModeler v2.0.0 for a comprehensive gene prediction set (45, 46). This gene set was also subjected to PASA expansion to identify additional splice variants. SAM file conversion was performed using SAMtools v1.16.1(47).

### Phylogenetic analysis

To identify and analyze the expression patterns of *DMRT* genes, we first performed an exhaustive search for genes harboring Doublesex-Mab-3 related domains (DM domains). Initial candidates were extracted based on the InterProScan searches performed above, and the corresponding amino acid sequences were submitted to a BLASTP v2.2.22 (48) search against *P. metropolitanus*. In addition, the amino acid sequences of *P. metropolitanus DMRT* orthologs were subjected to a BLASTP search against amino acid sequences predicted from various tardigrade genomes (32). The amino acid sequences for the tardigrade *DMRT* orthologs, metazoan orthologs provided in a previous study (49), and velvet worm *Dmrt* ortholog were pooled and then aligned with MAFFT v7.450 (50) and subjected to phylogenetic tree construction using IQTREE2 v2.2.2.6 (51). The phylogenetic tree was visualized in FigTree v.1.4.3 (http://tree.bio.ed.ac.uk/software/figtree). The expression patterns of *H. exemplaris* and *R. varieornatus DMRT* orthologs during developmental stages were obtained from our previous report (52). Additional alignments for the DMRT proteins were performed by MAFFT v7.450 and visualized using MView (https://www.ebi.ac.uk/Tools/msa/mview/).

Similarly, we conducted a phylogenetic analysis of *CAHS* genes. We obtained annotated CAHS sequences from our previous report (53) and pooled the amino acid sequences of *P. metropolitanus* CAHS candidate orthologs. A phylogenetic tree was constructed using the same procedure.

### Genome extraction

Virgin *P. metropolitanus* was prepared by the method described above, and a single tardigrade was placed in a 0.2 ml tube after 1% penicillin/streptomycin treatment for 2h to remove contamination. Genomic DNA was extracted and prepared using the method described by Arakawa et al. (25). An individual was crushed by pressing it against a tube wall using a pipette tip. Genomic DNA was extracted with Quick-gDNA MicroPrep kit (Zymo Research) with three freeze-thaw cycles and then following the manufacturer’s protocol. The extracted DNA was sheared to 550 bp target fragments with Covaris M220 and a Illumina library was constructed with a Thruplex DNA-Seq kit (Takara BioRubicon Genomics). Quantification, quality, and library size selection were performed with Qubit Fluorometer (Life Technologies) and TapeStation D1000 ScreenTape (Agilent Technologies), respectively. Sequencing library fragments in the range of 400–1,000 bp were cut and purified with a NucleoSpin Gel and PCR Clean-up kit (Clontech) and sequenced using a NextSeq500 sequencer with HighOutputMode 75 cycles kit (Illumina). The reads were de-multiplexed, and adaptor sequences were removed using the bcl2fastq v2 software (Illumina).

### Genome reassembly

Previously published ONT raw reads were submitted for reassembly using Canu v2.2 (54), NextDenovo v2.5.2 (55), Shasta v0.11.1 (56), Flye v2.9.2-b1786 (57), redbean v2.5 (wtgbt2) (58), GoldRush v1.1.0 (59), SPADES v3.15.5 (60), Pecat v0.0.3(61), and Raven v1.8.3 (62). Polishing was performed using NextPolish v1.4.1 (63) for the NextDenovo assembly. Each assembly was evaluated using compleasm v0.2.2 (metazoa and eukaryota lineage) or BUSCO v5.5.0 (64, 65). Completeness was also evaluated for *H. exemplaris* and *R. varieornatus* published genomes as well (31). The coverage for 10 kbp bins was calculated as previously stated, where we used BWA-MEM2 v2.2.1 (66) instead of BWA-MEM. *DMRT* orthologs were searched with TBLASTN v2.2.22, using *P. metropolinatus* DMRT protein sequences as the query (E-value < 1e-50). Additionally, we co-assembled male and female short reads produced in this study along with the ONT and Illumina datasets using SPADES v3.15.5.

### Sex specific region analysis

To identify candidate sex chromosome regions, we employed the Y chromosome genome scan (YGS) method (67), which was previously used to identify *Drosophila melanogaster* sex chromosome contigs. Briefly, reads from the same sex were pooled, and 15-mers were extracted with jellyfish count v2.2.4 or v2.2.10 (68). Scripts from the YGS method v.11b (8 Oct 2012 10AM) were then used to calculate the percentage of validated single-copy unique *k*-mers (P_VSC_UK) for each contig. This was performed for the previously published genome, as well as for the SPADES assembly performed above. We also tested the coverage for both sexes calculated from the gDNA-Seq data. The raw gDNA-Seq reads were mapped to the genome using BWA-mem2 v2.2.1 and converted into BAM files using SAMtools v1.17. The genome was split into 10 kbp bins and the average coverage for each bin was calculated using BEDtools v2.31.0. The values were then normalized by the median of all bins for that sample, and the average for males and females was computed.

For gene-level synteny analysis, we employed the Python version of McScan in the JCVI suite v1.2.7 (69). Gene prediction and coding sequences were prepared for *H. exemplaris* and *P. metropolitanus* and syntenic regions were identified and visualized using default settings. To identify the *Dsup* ortholog, candidates were identified using gene-level synteny. Disorderness was analyzed using DISOPRED (http://bioinf.cs.ucl.ac.uk/psipred/) and IUPRED3 (https://iupred3.elte.hu/) and the protein structure predicted ColabFold v.1.5.3 (34, 70, 71).

### Genotyping for male specific regions

Virgin specimens were replaced with single-worm lysis buffer (50 mM KCl, 10 mM Tris pH8.2, 2.5 mM MgCl2, 0.45% NP-40, 0.45% Tween20, 0.01% gelatin, 2 μg of Proteinase K) (72). The specimen was then dissolved by freeze-thaw cycles (three times for liquid N2 and RT) and incubated at 60℃ for 1.5 h and 95 °C for 25 min. Genotyping PCR was performed using the following conditions: 94 °C for 3 min; 40 cycles of 94 °C for 30 s, 50 °C for 30 s, and 68 °C for 1 min; and a final extension at 68 °C for 5 min. Primer sequences were designed using Primer3 (73) from the nucleotide sequences of scaffold Parri_scaffold0000295 (**Supplementary Table S1)**. Quick-Taq (TOYOBO) was used for the polymerase with the concentrations of each reagent, following the manufacturer’s instructions. Electrophoresis was performed at 100V for 20 min with 1.2% agarose gel/TAE (NacalaiTesque), and then the gel was strained with ethidium bromide for 20 min. The DNA bands were visualized using ChemiDoc (BioRad).

## Results

### Transcriptomics of P. metropolitanus sexes

To identify sex-specific gene expression and genomic loci, we produced 10–20 M reads of RNA-Seq data for male and female specimens (**Supplementary Table S2**) that mapped approximately 80–90% of the genome. Based on these data, we quantified and conducted differential gene expression analysis. PCA analysis of the expression profiles indicated a clear distinction between the male and female samples (**Figure 1A**). A total of 9,015 transcripts were differentially expressed, with 4,685 and 4,329 transcripts showing higher expression in females and males, respectively (**Figure 1B**). Gene ontology enrichment analysis of each gene set indicated enrichment of various pathways (**Supplementary Figure S1**). For females, we observed enrichment of RNA processing, cellular component biogenesis, and negative regulation of biological processes. In contrast, terms related to cyclic nucleotide biosynthetic processes, aminoglycan metabolic processes, and monatomic ion transport were enriched.

**Figure 1.**
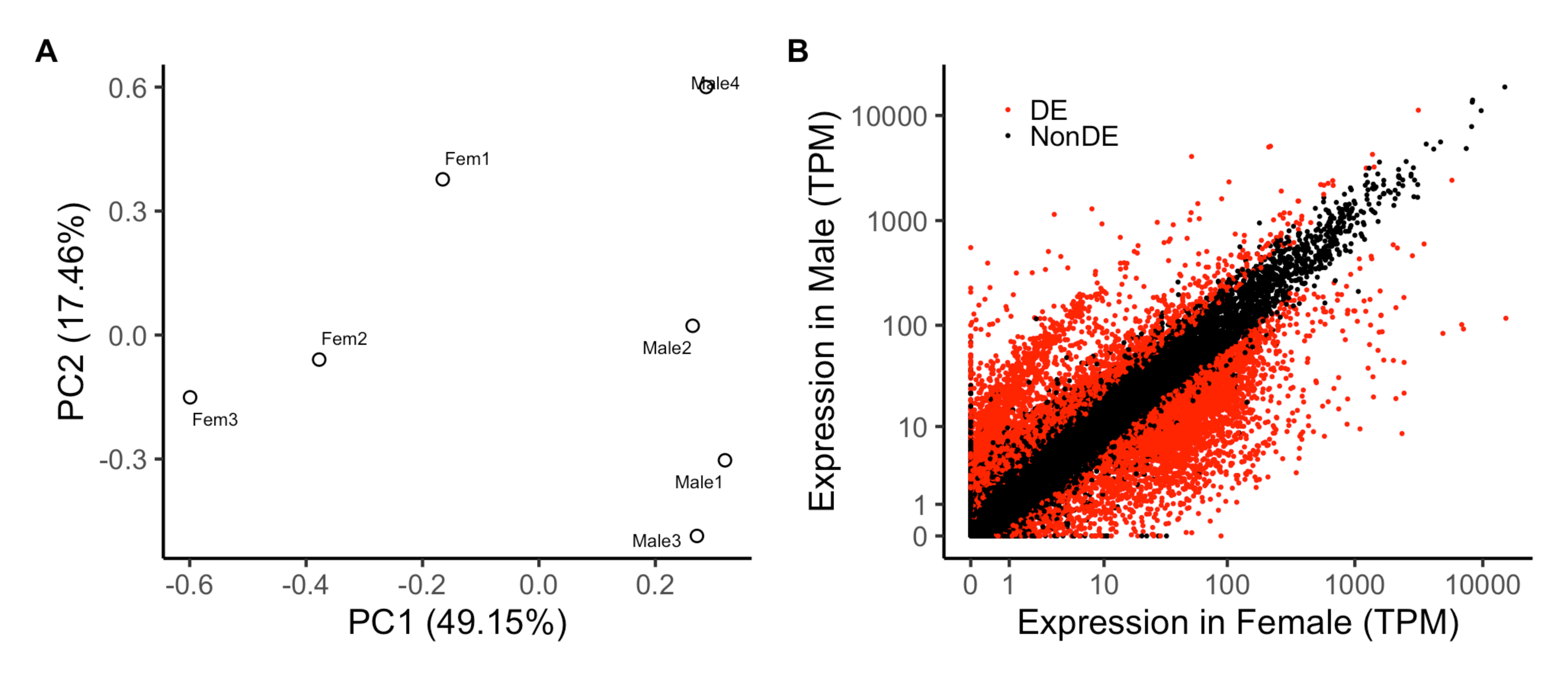
Transcriptomic analysis of both sex. **[A]** PCA analysis of expression profiles. **[B]** Scatterplot of the expression profiles. Red dots indicate differentially expressed transcripts (FDR<0.05).

The sex determination cascade comprises multiple genes, forming a signaling cascade that causes differentiation between the sexes. We first focused on *DMRT*, a well-conserved gene family that regulates sex-specific tissue development and behavior. Initial BLAST analysis identified five *DMRT* orthologs (PARRI_0009851, PARRI_0001169, PARRI_0005877, PARRI_0003090, and PARRI_0003093). We observed that three genes, PARRI_0003090, PARRI_0003090, and PARRI_0005877, were upregulated in males, whereas PARRI_0003090 was moderately expressed (TPM>30) in males. Although we could not determine *sxl* and *fru* orthologs, we identified possible sex specific variants for the *tra2* ortholog (**Supplementary Figure S2**).

A diverse array of lineage-specific upstream signaling factors (e.g., *tra2*, *nix*, *fem*) induce sex-specific splicing variants of the *doublesex* gene, transmitting signals to the downstream sex development cascade (6). Although the master regulator of sex determination is highly variable, several components of the cascade are highly conserved, such as the *DMRT* orthologs. Likewise, the *Transformer-2* (*Tra2*) gene is a DNA-binding protein coupled with the Tra protein, causing sex-specific splicing of *dsx* in insects (6). The *P. metropolitanus Tra2* (*PmTra2*; PARRI_0000692) exhibited sex-specific variants with different splicing sites in the second exon (**Supplementary Figure S2**). The female variant of *PmTra2* (*PmTra2F*, evm.model.Parri_scaffold0000002.194) has an extra transcribed region at the 5’ end, whereas the start site for the male variant (*PmTra2M*, evm.model.Parri_scaffold0000002.194.3.65434fff) is located on the third exon.

We also attempted to identify genes participating in the sex cascade in *Drosophila*, e.g., *fruitless* (*fru*) and *sexlethal* (*sxl*) genes. BLAST searches identified two candidates for *sxl* orthologs (PARRI_0002227 and PARRI_0007430) which showed contrasting expression profiles. However, a phylogenetic analysis indicated that these genes could not determine whether these orthologs were *sxl* or a gene family with relatively high similarity; therefore, we cannot conclude whether these genes are *sxl* orthologs (data not shown). No hits were found for *fru* in *P. metropolitanus* nor any tardigrade genomes. Thus, we concluded that *fru* is missing and *sxl* remains questionable in *P. metropolitanus*.

Taken together, we conclude that sex-specific *tra2* and *DMRT* exist and may be functional in the *P. metropolitanus* sex determination cascade; however, several factors of this cascade may be lost in this lineage. The *Bombyx mori sxl* gene induces dimorphism of the sperm, not sex determination (74); therefore, it is possible that the lack of *sxl* may imply a different regulatory pathway than is known.

### Genomic loci of the male-biased genes

We detected a peculiar population of genes that were approximately >25 higher expressed in males (**Figure 1B**). Hypothesizing that these male-biased expressed transcripts may be sex-specific genes located on the sex chromosome, we conducted a genomic enrichment analysis to determine genomic loci enriched in these highly biased genes. Using a genomic bin of 200 kbp (corresponding to roughly 30 genes per bin) against differentially expressed transcripts that had over 10x fold change then the other sex (Female: 674, Male: 1724), we detected 325 (29 scaffolds) and 12 (3 scaffolds) bins for males and females, respectively (**Figure 2A**). We noted that approximately 2% of the male-biased genes had more than TPM 10 in females (11% for male-expression of female-biased genes), thus implying the specificity of male-biased genes. Gene ontology enrichment analysis of genes located in these bins indicated a high enrichment of transcripts related to sperm function (**Supplementary Table S3, S4**). Interestingly, two out of the three male-induced *DMRT* paralogs (PARRI_0003090 and PARRI_0003093) were located within a bin enriched for male-biased genes on the scaffold Parri_scaffold0000005 (**Figure 2B**). We also observed that the genes within and in the surrounding regions of these bins were also expressed in females, suggesting that these genomic loci may not be male-specific (**Figure 2C**).

**Figure 2.**
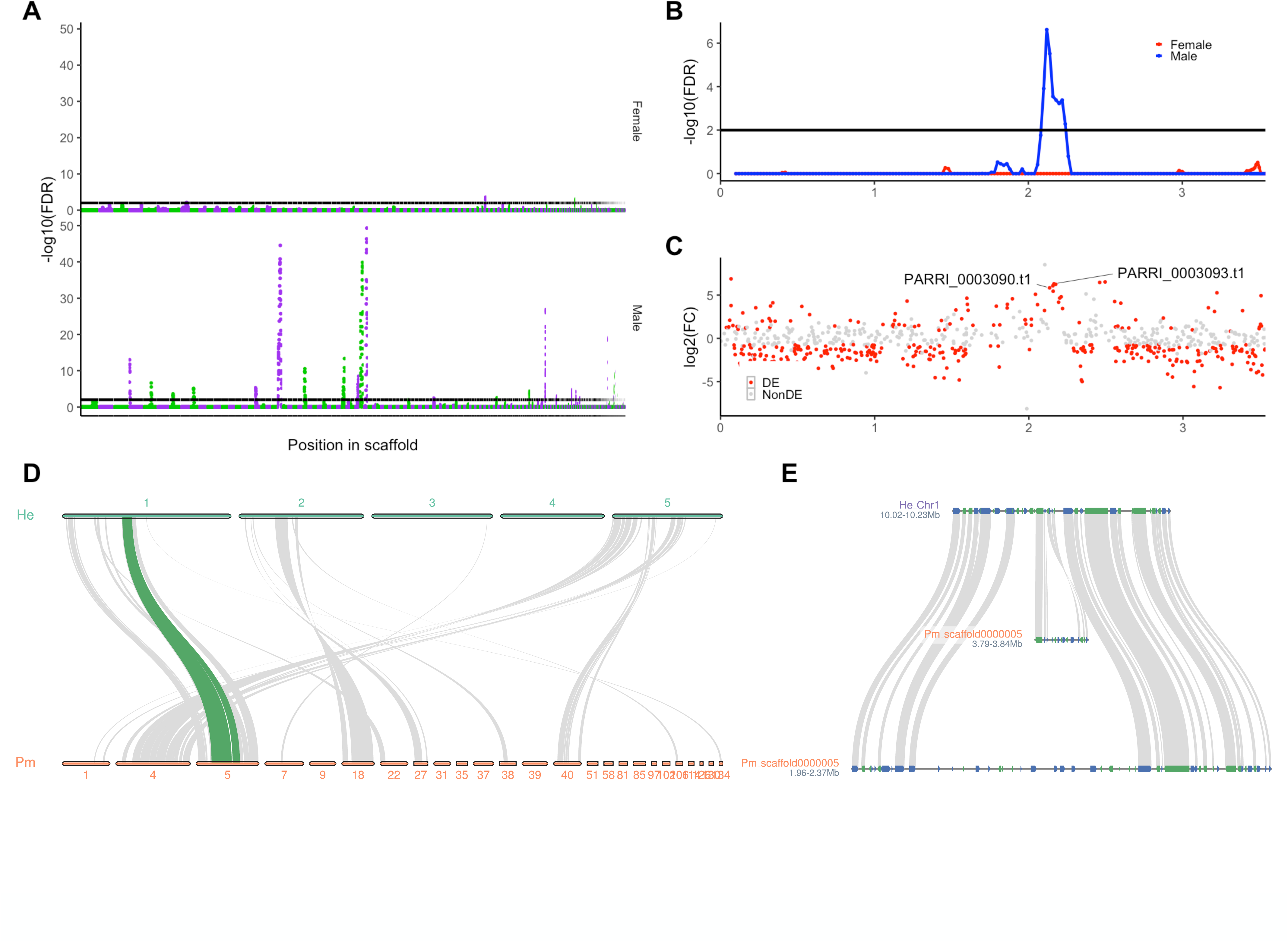
Multiple male-biased regions within the *P. metropolitanus* genome and their synteny. **[A]** Genome-wide enrichment analysis of male- or female-biased transcripts. Scaffolds were ordered by size and colored green and purple to visualize the scaffolds. The threshold of FDR < 0.01 was used. **[BC]** Characteristics of scaffold Parri_scaffold0000005 harboring the *DMRT* paralogs. **[B]** FDR values from the genomic loci enrichment analysis plotted against the bin’s position. Blue and red indicate the values for males and females, respectively. **[C]** Expression fold-change log2(male + 0.1 / female + 0.1) of the genes plotted against their location along the scaffold. Colors indicate whether the gene was differentially expressed. **[D]** Macro-scale synteny analysis to identify orthologous genomic loci in male-biased scaffolds. Gray lines indicate syntenic blocks between *H. exemplaris* and *P. metropolitanus* scaffolds. The synteny block highlighted in green indicates the location of *DMRT* paralog loci. The numbers on the bar indicate the chromosome number or scaffold ID for each genome assembly. **[E]** Synteny region of the *DMRT* paralog loci on *P. metropolitanus* scaffold Parri_scaffold0000005 and *H. exemplaris* Chromosome 1.

To evaluate whether these genomic loci enriched for male-biased genes were on the same chromosome, we preformed synteny analysis with the recently reported chromosome-level genome assembly of *H. exemplaris*. *Paramacrobiotus metropolitanus* has been observed to have 2n=10 karyotypes, similar to that of *H. exemplaris* (21, 75). While the queried 29 male-biased scaffolds did not focus on a particular chromosome, we observed a slight bias toward chromosomes 1, 2, and 5 (**Figure 2D**). Furthermore, we observed that the paralogous *DMRT* loci and the surrounding region on Parri_scaffold0000005 were missing in *H. exemplaris*, and different loci on the same chromosome were inserted into the corresponding region (**Figure 2E**). These data suggest that this genomic region may have emerged in the *P. metropolitanus* lineage.

### Emergence of a novel dmrt93B-like subfamily specific to Macrobiotidae

Considering the importance of paralogous *DMRT* genes located on Parri_scaffold0000005, we focused on the characterization of the orthologs to determine the characteristics of these paralogs.

First, we submitted the amino acid sequences of the *P. metropolitanus* DMRT family for phylogenetic analysis, incorporating various tardigrade *DMRT* orthologs from genome and transcriptome assemblies. Careful examination of the *PARRI_0003093* gene structure revealed a misassembly of a single nucleotide insertion, identified through gDNA- and RNA-Seq read mapping. This caused a frameshift in the 3’ terminus, leading to a truncated coding sequence. Therefore, manual curation for this gene was performed, resulting in 463 amino acid sequence. Phylogenetic analysis identified *PmDmrt99B* (PARRI_0009851), *PmDmrt93B* (PARRI_0005877), and *PmDmrt11E* (PARRI_0001169) orthologs, as well as two *Dmrt93B*-like paralogs (*PmDmrt3090* PARRI_0003090; *PmDmrt3093* PARRI_0003093; the 3090/3093 complex). The 3090/3093 complex contained *DMRT* genes only from Macrobiotidae species, suggesting the acquisition of this subfamily in this lineage (**Figure 3A**). We also observed a phylum-wide loss of the *Doublsex* subfamily. Furthermore, we observed an Echiniscidae-specific Dmrt93B subfamily that was not included in the 3090/3093 complex. While the relatively lower bootstrap support of this branch (88) complicated the phylogenetic position of this clade, only the DM domain was found in these subfamily members. Interestingly, phylogenetic analysis indicated that the *dsx*-like gene of the velvet worm branched into a Doublesex clade with arthropods, suggesting that *dsx* emerged after the divergence of Tardigrada. We were unable to detect two copies of the 3090/3093 complex in several other Macrobiotidae species. A direct comparison between PmDmrt3090 and PmDmrt3093 amino acid sequences indicated that the first 30–180 aa sequences were extremely similar, but the intron nucleotide sequences were completely different. Furthermore, multiple nanopore reads spanned the entire length of each gene. Together, we suggest that the two copies were not the result of misassembly of these loci. The lack of two copies in other Macrobiotidae species may be the result of misassembly in their genomes; the analyzed genomes are based on Illumina short reads, and the extremely similar 30–180 aa (corresponding to approximately 450 bp) may have resulted in a misassembly. We also noted that no ONT reads spanned both *PmDmrt3090* and *PmDmrt3093*. However, the 3090/3093 complex region spanned for more than 30 kbp and the N50 length of the ONT data was approximately 17 kbp. It is possible that there are not ONT reads spanning the entire region. While reassembly of the ONT reads using more recent assembly methods produced a more contiguous assembly (NextDenovo + NextPolish; **Supplementary Table S5**), these two genes were predicted to be two separate genes.

**Figure 3.**
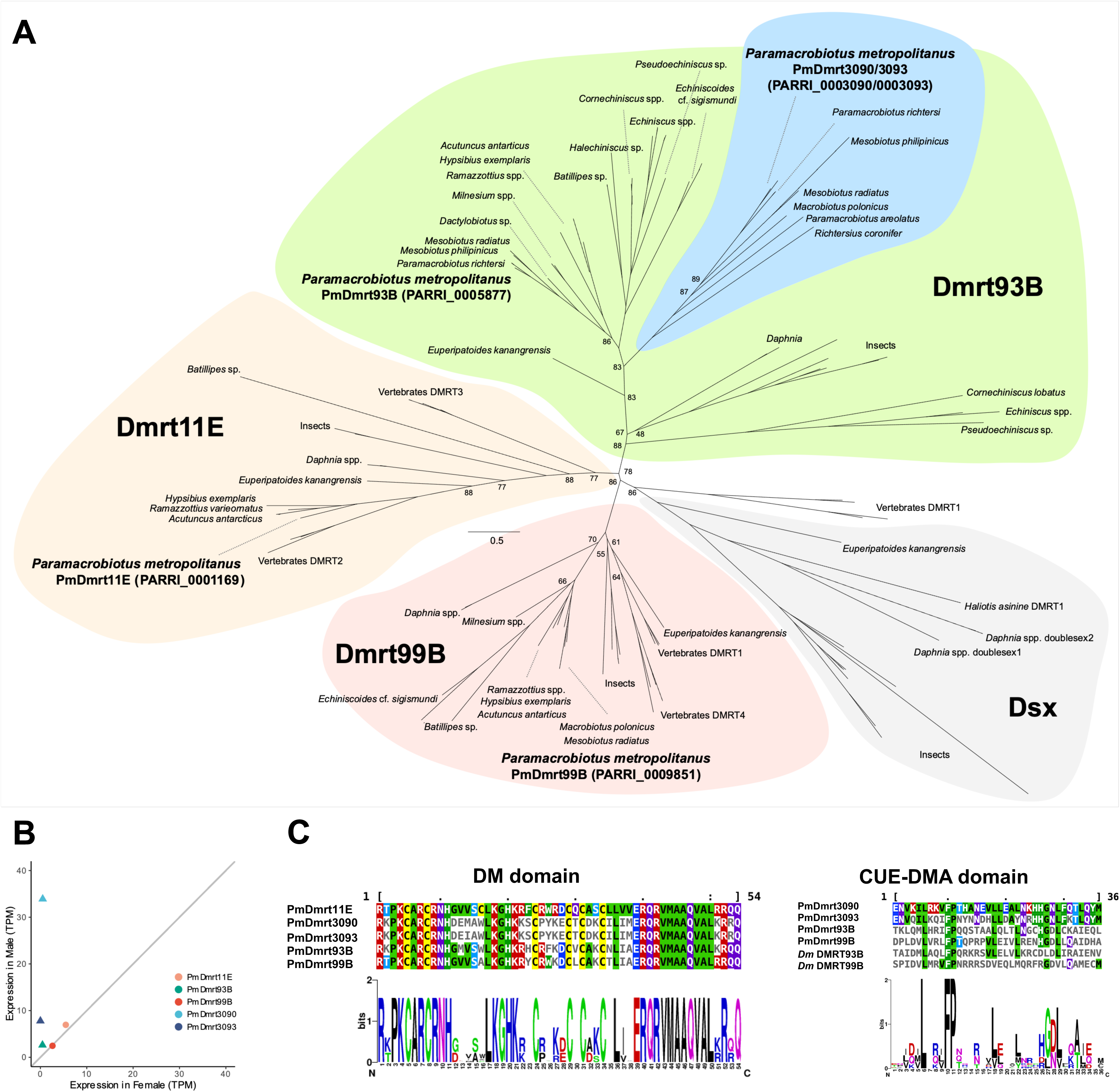
Phylogenetic analysis and the expression of *DMRT* orthologs. **[A]** Phylogenetic analysis of DMRT orthologs detected in the tardigrade genomes. The DMRT families were classified based on the orthologs of the model species. Bootstrap values of less than 90% are shown on the branch. **[B]** Expression of *PmDmrt* orthologs. Triangle points indicate differentially expressed genes and circles indicate non-significant changes. The gray line indicates x=y. **[C]** Multiple alignments of DM and CUE-DMA domains. *Dm* indicates *D. melanogaster*.

Based on these annotations, we identified *PmDmrt3090*, *PmDmrt3093*, and *PmDmrt93B* to be significantly expressed in males; thus, all three induced copies belong to the Dmrt93B clade (**Figure 3B**). To evaluate the expression of *DMRT* orthologs in other tardigrades, we utilized our previously reported single specimen RNA-Seq data of the embryonic and juvenile life stages of the parthenogenetic tardigrades *H. exemplaris* and *R. varieornatus*. Only females have been observed in both species, suggesting the lack of masculinization in these species. All three *Dmrt11E*, *Dmrt93B*, and *Dmrt99B* orthologs in *H. exemplaris* and *R. varieornatus* (RvDmrt11E: g5527, RvDmrt93B: g9000, RvDmrt99B: g7078; HeDmrt11E: BV898_08851, HeDmrt93B: BV898_13063, HeDmrt99B: BV898_01934.) were expressed during embryonic stages (**Supplementary Figure S3,**), where *Dmrt11E* preceded *Dmrt99B* in both species, and the three *DMRT* genes were expressed at lower levels in juvenile and adult stages.

We further investigated the functionality of the *DMRT* orthologs by functional domain detection (**Figure 3C**). While all five DMRT copies harbored the DM domain at the N-terminus, they did not contain the dimerization domain known to exist in *dsx* proteins required for DNA binding and sex-specific splicing variants. While we detected the DM domain in all five orthologs, we did not find a ubiquitin biding-related CUE-DMA domain only in PmDmrt3093 but not in PmDmrt3090 by sequence-based domain search analysis. Multiple alignment of the five orthologs and the *D. melanogaster* DMRT sequence suggested the conservation of several residues within the region corresponding to the CUE-DMA domain, implying the conservation of this domain. By modeling the protein structure with AlphaFold2 and aligning the *D. melanogaster* Dmrt93B structure, we observed that the C-terminal region showed structural homology with CUE-DMA domain-like helices (RMSD: 0.276–0.574, **Supplementary Figure S4**), suggesting that PmDmrt3090 may also harbor the CUE-DMA domain. These data and the lack of the *dsx* subfamily suggests that the sex determination cascade may differ from that of the dsx paradigm in insects, by utilizing the 3090/3093 complex paralogs(76).

### Contradicting data between whole genome sequencing and PCR based genotyping

Based on our observations of several male-biased but not male-specific genomic regions, we hypothesized that these regions were not sex-specific chromosome structures. To evaluate this, we sequenced the genomes of both sexes at low coverage. We produced approximately 50–60M reads, corresponding to roughly 20–25x coverage (**Supplementary Table S1**). Approximately 80–90% of the reads were mapped to the genome, resulting in roughly 15–20x coverage.

We first calculated the coverage of the 10 kbp bins genome wide. Initial PCA of the coverage profiles did not show a clear difference between males and females (**Figure 4A**). We identified several bins with half of the average genome-wide coverage that were not found in females (**Figure 4B**). These characteristics are similar to those of heterozygotic chromosomes, particularly the X chromosome of males in the XY sex determination system. All of the bins that were identified as male-biased by the transcriptome analysis had genome-wide average coverage, suggesting that all regions exist in females (**Figure 4C**). We also evaluated whether we could detect male or female specific regions through *k*-mer based analysis using the YGS method. Scaffolds that have a high number of “percent validated single-copy unmatched *k*-mers” (P_VSC_UK) indicate sex specificity. Although no scaffolds had a P_VSC_UK ratio of 100, we detected five scaffolds fitting the XY sex chromosome structure rather than the ZW scheme with an arbitrary threshold of P_VSC_UK > 80 (**Figure 4D, E, F**). Similar profiles were observed by SPADES reassembly using all gDNA-Seq data from our and previous studies (**Figure 4G, H, I**). We also noticed that many scaffolds from both assemblies had P_VSC_UK values of approximately 50% in both female-to-male (XY) and male-to-female (ZW) analyses (**Figure 4F, I**), which indicates that the corresponding region is both male- and female-specific.

**Figure 4.**
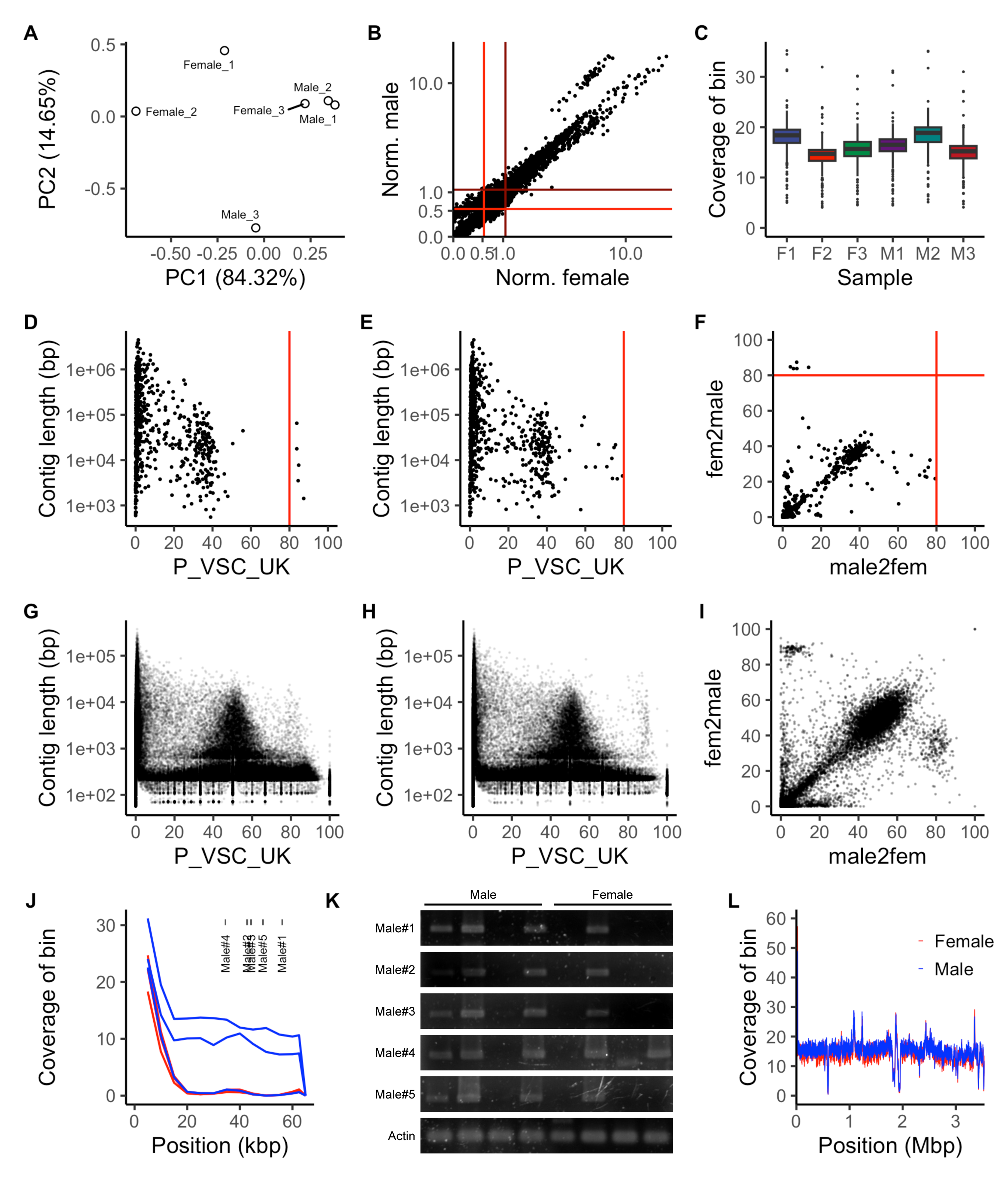
*P. metropolitanus* lacks sex specific regions for both sex. **[A]** PCA of genomic coverage profiles for male and female gDNA-Seq data. **[B]** Average coverage for 10kbp bins, normalized by the median of all bins. The brown and red lines indicate the whole-genome average and half-genome average for males and females, respectively. The range between the 0x and 10x coverage ratio to the median is shown. **[C]** Genome coverage of male-biased bins. **[DEGH]** YGS analysis for **[DG**] female-to-male (XY system) and **[EH]** male-to-female (ZW system) based on the **[DE]** published *P. metropolitanus* assembly and **[GH]** SPADES reassembly. The red line indicates a P-VSC-UK threshold of 80. **[FI]** Scatter plot for P-VSC-UK values for male-to-female and female-to-male analysis for the **[F]** published genome and the **[I]** SPADES reassembly. Contigs shorter than 1,000 bp were removed from the SPADES plot. **[J]** gDNA-Seq coverage for all samples and the location of genotyping primers within contig Parri_scaffold0000295. Blue and red correspond to male and female samples, respectively. **[K]** Electrophoresis of genotyping primers designed for male specificity. Male specificity was not observed for any primer set. **[L]** gDNA-Seq coverage of *PmDmrt3090*/*PmDmrt3093* harboring scaffold Parri_scaffold0000005 for all samples. Colors indicate samples for each sex.

To evaluate the male specificity observed in the *in-silico* analysis, we designed several primers to amplify regions in the scaffold Parri_scaffold0000295 that were identified as male-specific (**Figure 4J, Supplementary Table S1**). Evaluating individual genomic coverage indicated that in this scaffold, a single male sample had near-zero coverage, in contradiction with the other two male samples (**Figure 4J**). However, PCR genotyping indicated the existence of this region in females as well, which contradicted the results obtained from the *in-silico* analysis (**Figure 4K**). Thus, we concluded that we could not derive sex chromosomes or male-specific regions, and the male-specific regions detected above may have been an artifact of differences between individuals. We also evaluated the male-specificity of the paralogous DMRT loci on Parri_scaffold0000005, where coverage analysis suggested that this region was not male-specific (**Figure 4L**).

We noticed relatively low level of RNA-Seq mappability to the reported genome (∼90%), which lead us to re-evaluate the current genome. Completeness analysis indicated 72.9% completeness which the most recent version of BUSCO. Furthermore, we observed several scaffolds with inconsistent coverage distribution in our sex-separated data, but not in Hara *et al.* Illumina data. Therefore, we tested if recent assemblers would result in a more contiguous and complete assembly, compared to the Canu assembled current genome. However, we were not able to obtain a more complete genome, with the maximum being a 0.4% increase for the assembly derived with NextDenovo + NextPolish (**Supplementary Table S5**). Other statistics had a large increase; N50 from 1.0M to 1.3M, longest scaffold length 4.48M to 9.23M. For comparison, we evaluated completeness of other high-quality tardigrades genomes, namely *R. varieornatus* and *H. exemplaris*. Both BUSCO and compleasm resulted in completeness values similar to *P. metropolitanus*; *R. varieornatus* (C:74.6%) and *H. exemplaris* (C:73.3%). These data suggests either tardigrade genomes may lack some BUSCO genes, or the gene detection algorithm of the current BUSCO software may not fit the genome of tardigrades, resulting in lower BUSCO scores. Therefore, we used the current genome for *P. metropolitanus* for later analysis.

### Sex-bias in anhydrobiosis related genes

A major feature of tardigrades is their ability to survive the extremities, a phenomenon known as cryptobiosis (77). Tolerance to near-complete desiccation is known as anhydrobiosis (78). Several tardigrade-specific gene families, *i.e.* cytosolic-abundant heat soluble (CAHS) and Secretory Abundant Heat Soluble (SAHS), have been implicated in anhydrobiosis protection (20). A recent study observed tissue-specific expression of anhydrobiosis genes (79). Both males and females are capable of anhydrobiosis, in which protective genes are expressed in sex-specific organs, such as the testes or ovaries. Therefore, we hypothesized the presence of sex-biased anhydrobiosis genes.

We used our previously reported RNA-Seq data for the hydrated active state and the tun state, desiccated for two days, to identify genes induced during anhydrobiosis. We detected approximately 4,500 differentially expressed transcripts, slightly fewer than in our previous report, possibly due to the different methods used for differential expression analysis. We then compared the expression profiles of anhydrobiosis and between sexes and observed approximately 1,800 transcripts that were differentially expressed under both conditions (**Figure 5A**). As hypothesized, we observed that three CAHS and one SAHS ortholog were sex-biased, possibly implying tissue specificity (**Figure 5A**). Interestingly, all three CAHS orthologs induced in males were the only three among the 13 CAHS orthologs that were not differentially expressed during anhydrobiosis (**Supplementary Table S6**). Phylogenetic analysis indicated that these CAHS orthologs were CAHS1 (PARRI_0016931), putative CAHS5 (PARRI_0006576), and CAHS5 (PARRI_0002229) orthologs, following the proposed naming scheme of Fleming *et al.* (53). In contrast, the SAHS ortholog, detected as differentially expressed, was induced in the females. We also found six orthologs of tardigrade-specific manganese-dependent peroxidase (32) to be highly expressed in males but not in females. Only four genes were found to be induced during anhydrobiosis.

**Figure 5.**
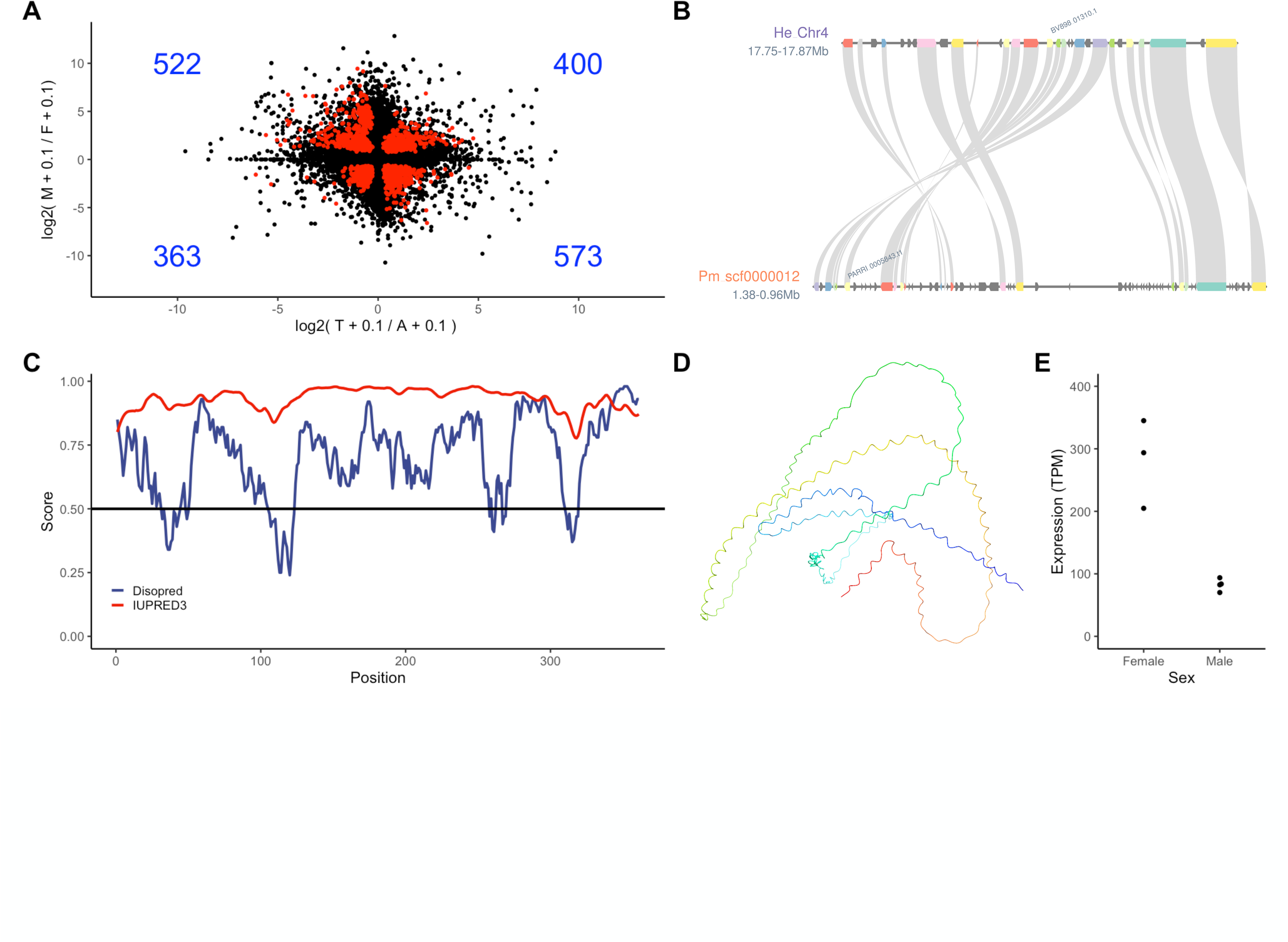
Sexual bias in anhydrobiosis genes and identification of PmDsup ortholog. **[A]** Comparison of gene expression profiles between the sexes during anhydrobiosis. Log2 (Tun +0.1) / (Active + 0.1) were plotted for the x-axis and for the y-axis log2 (Male + 0.1) / (Female + 0.1). Red dots indicate transcripts detected as differentially expressed in both comparisons and the number in each quadrant is indicated in blue text. **[B]** Synteny analysis to identify orthologous genomic loci in *H. exemplaris* and *P. metropolitanus*. **[C]** Disopred and IUPRED3 scores **[D]** Protein structure predicted by ColabFold. The N-terminus to the C-terminus shows gradient colors from blue to red. **[E]** Expression of *PmDsup* in both sexes.

Based on the identification of the *H. exemplaris* ortholog of the *Damage suppressor* (*Dsup*, BV898_01301) gene, we also searched for a *P. metropolitanus Dsup* ortholog through gene synteny with *H. exemplaris* (30, 80). We identified PARRI_0005796 as a *Dsup* ortholog candidate (**Figure 5B**). This protein was annotated as “transcriptional regulatory protein AlgP’’ in NCBI; however, (1) no functional domains were identified by InterProScan, (2) no BLAST hits to known proteins (E-value < 1e-5), (3) highly disordered throughout the whole protein (**Figure 5C**), and (4) a predicted nuclear localization signal (DeepLoc2, 0.7715 probability), suggesting that this protein may be a *Dsup* ortholog. The AlphaFold2 structure prediction also implied a lack of globular structure (**Figure 5D**). *PmDsup* was significantly upregulated in females (TPM, female: 280, male: 82, FDR = 1.31 x 10^-6^, **Figure 5E**), implying the importance of this gene in females.

## Discussion

In this study, we focused on gonochoristic tardigrade *P. metropolitanus* to identify possible factors that affect sexual dimorphism. Cytological studies have not identified definitive sex-linked chromosomes in tardigrades (81, 82) and multiple reports have observed biased sex ratios in tardigrades (21, 83-86). These observations suggests that the sex determination of tardigrades may not depend on the random distribution of sex chromosomes (or the existence of a sex chromosome). Even in the absence of sex chromosomes, as hypothesized in tardigrades, genomic loci affecting sexual dimorphism would exist, which may be detected by comprehensive omics methods.

Therefore, we aimed to characterize the molecular basis of sexual dimorphism in tardigrades by comparing the transcriptome and the genome between *P. metropolitanus* sex. We hypothesized that sex-linked genes may be related to sex determination or dimorphism, and if focused on a small genomic region, may imply a sex-determining region, such as the M factor found in many eukaryotes (87). Transcriptome analysis of both sexes indicated a large number of sex-biased genes, despite the small morphological sex-linked differences in Macrobiotidae, with the exception of their germline (18). We observed upregulation of genes related to spermatogenesis in males, which reflects the activation of spermatogenesis, and large amounts of sperm are continuously produced in adult males (21, 84). In contrast to that in males, DNA replication- and meiosis-related genes were highly expressed in females. Females undergo DNA replication not only to produce oocytes through meiosis (21) but also to shed the cuticular shell during the last stage of the reproductive process (simplex stage) (21). Mitotic cells are generally observed in the post-simplex stage (88). Together, the regulation of DNA replication and meiosis is consistent with the production of mitotic cells and extensive replication of the epidermal layer (88, 89).

We identified a small gene set highly biased toward males but missing in females, which we hypothesized may be related to sexual dimorphism. Genome loci enrichment analysis of this gene set identified approximately 325 bins spanning 29 scaffolds as male-biased. This region was enriched in sperm and ion transport-related genes, which is consistent with the production of sperm at the adult male life stage. To evaluate sex specificity, we produced low-coverage genome sequencing data to evaluate sex-specific regions and observed that most regions were present in the genomes of both sexes. Genome-wide analysis revealed several male-specific regions; however, PCR evaluation produced contradictory results. We used a laboratory-cultured TYO strain of *P. metropolitanus* for genome and transcriptome sequencing, therefore, we anticipated low levels of heterozygosity within the culture population. However, the results obtained at this stage implied that the genomic differences we detected as sex-linked can be explained as individual variability. Additionally, during the YGS analysis, we observed a high number of contigs that showed approximately 50% P_VSC_UK, suggesting that there are a large number of contigs that contain sequences specific for both sexes, which we hypothesize that individual variability may have caused this abnormal distribution. Together, the lack of sex specific regions may imply that the difference between sexes may be due to epigenetic modifications.

One of the key findings of this study is the accumulation of knowledge for sex determination cascade-related genes, particularly the *DMRT* gene family. The DMRT family is a highly conserved transcription factor that plays an important role in sex differentiation in many animals and has been studied extensively in insects (7). Several studies have identified *DMRT* orthologs to be located on the sex chromosomes and regulate the growth of sex-specific tissues (8, 9). The evolutionary background of this gene family has been extensively analyzed in other lineages (7); however, such analysis has been overlooked. In our analysis, we identified a Macrobiotidae-specific DMRT93B subfamily located in a male-biased region, which we termed the 3090/3093 complex in addition to the *Dmrt99E*, *Dmrt93B*, and *Drmt11E* subfamilies. Orthologs of this subfamily have been found in Macrobiotidae and several Echiniscidae. While conservation in Echiniscidae complicates the evolution of this subfamily, the identification of orthologs in various Macrobiotidae species suggests that this is an important *DMRT* subfamily. In fact, the two 3090/3093 complex paralogs were expressed higher in males, similar to *Daphnia Dsx1* (11, 12), suggesting that these subfamily orthologs may inhibit feminization or progress musculation. Furthermore, we did not find any orthologs of the *dsx* subfamily in any of the tardigrade genomes analyzed, and did not identify splicing variants in any of the *P. metropolitanus DMRT* orthologs, suggesting a sex differentiation cascade different from those that rely on sex-specific *dsx* splicing variants like those observed in insects.

Tardigrades are renowned for their ability to tolerate extreme stress (20), and *P. metropolitanus* also shows a high tolerance to desiccation stress (22). Interestingly, we observed sex-biased expression of several anhydrobiosis genes, hypothesized to play protective roles during anhydrobiosis (29-32, 90). For instance, CAHS genes are tardigrade-specific proteins that form gel filaments that possibly protect cells (91-93). Recent studies have observed tissue/organelle specificity for these proteins, which further implies the existence of orthologs with sex-specific expression (79). Therefore, we hypothesized that orthologs of such genes may exhibit sex-specific expression to protect sex-specific organs. Indeed, we identified CAHS, SAHS, and AMNP orthologs with sex-specific expression. Two of the three male-induced CAHS orthologs were highly expressed but were not induced between active and anhydrobiotic conditions. This may imply that these CAHS orthologs participate in the protection of male-specific tissues or sperm. Furthermore, we identified a *P. metropolitanus Dsup* ortholog that is highly expressed in females. Coupled with the observation of the enrichment of meiosis-related genes from transcriptome analysis, we suggest that *Dsup* may actively function to accommodate the production of oocytes/oogenesis rather than spermatozoa/spermatogenesis. In contrast, AMNP, a tardigrade-specific peroxidase, was highly expressed in males, suggesting enhanced protection against oxidative stress. Similar observations have been made in the sperm of many animals (94, 95). Together, the sex-biased expression of anhydrobiosis genes may provide protection for sex-specific tissues.

## Conclusions

In this study, we identified male-biased regions that may harbor potential candidates that regulate sexual dimorphism in the gonochoristic tardigrade *P. metropolitanus*. Simultaneously, these data denied the sex-chromosome-based sex determination scheme. We also provide evidence for a new *DMRT* subfamily that may contribute to sex differentiation in this family. The 3090/3093 complex DMRT paralogs may be initial candidates for disruption or gene editing for evaluation their relationships with sex determination (79, 96-98). Future studies utilizing high-quality genomes and careful physiological experiments are required to reveal sex determination cues not only in this species but also in other tardigrades.

## Declarations

### Ethics approval and consent to participate

Tardigrades (invertebrate) were used for this study. Any vertebrates, human, and their tissues were not applicable. The laws and regulations set forth by the Ethics Committees of Keio University and the University of Tokyo were followed.

### Consent for publication

Not applicable.

### Availability of data and materials

The raw reads for genome DNA-Seq were submitted to NCBI SRA under Bioproject PRJNA1063779. The raw reads and processed expression profiles were uploaded to NCBI GEO under the accession ID GSE253242. Other datasets analyzed in this study have been uploaded to figshare (10.6084/m9.figshare.25097525).

### Competing interests

The authors declare that they have no competing interests.

### Funding

Grant-in-aid KAKENHI (JP18J21345) to KS, (21H05279) to KA, grant for Research Project from Research and Education Center for Natural Science, Keio University for MM and KS. Joint Research of the 13 Exploratory Research Centers on Life and Living Systems (ExCELLS program 19-501, 22EXC601) to KA.

### Author contributions

KS, KH, and MM prepared specimens. KA performed the genome sequencing. KS, TK, and MM performed RNA-seq. KS and YY analyzed the data. KS and YY drafted the manuscript. AK, TK, and MM improved the manuscript. KS, YY, AK, TK, and MM designed this study.

## Acknowledgements

We thank Yuki Takai and Naoko Ishii (Keio University) for experimental support. For giving us many helpful comments, we also thank Dr. Hajime Watanabe (University of Osaka), Dr. Yasuhiko Kato (University of Osaka), Dr. Atsushi C. Suzuki (Keio University), Yu Saito (Keio University), and Ryo Ogushi (Keio University). RNA-seq was performed by Chemical Dojin Co. Ltd. We also thank the members of the Japanese Society for Tardigradology for fruitful discussions.

## Figure legends

**Supplementary Figure S1. Gene ontology enrichment analysis of differentially expressed genes between females and males.**

Gene ontology terms enriched in genes higher expressed in **[A]** females and **[B]** males.

**Supplementary Figure S2. *Tra2* genes of *P. metropolitanus*.**

Gene structure and RNA-seq-based evidence of intronic regions. From the upper row, (1) the gene structures predicted by Hara *et al.*, (2–4) RNA-Seq read mapping of male samples, (5– 8) RNA-Seq read mapping of female samples, and (9) EvidenceModeler and PASA expanded gene structure. These structures were visualized using Jbrowse2 instance.

**Supplementary Figure S3. Expression of *DMRT* orthologs in *H. exemplaris* and *R. varieornatus*.**

Error bars indicate the standard deviation. On the X-axis, E and B time points indicate #day after oviposition (embryo) and #days after hatching (baby), and adults (active and tun).

**Supplementary Figure S4. Structures of *DMRT* orthologs**

AlphaFold2 predicted the 3D structure of **[A]** full-length **[B]** DM domain, and **[C]** the CUE-DMA domain. The arrowheads in cyan and magenta indicate the DM and CUE-DMA domains, respectively. *Dm* indicates *D. melanogaster*.

## Notes

### Competing Interest Statement

The authors have declared no competing interest.

