## Supplementary figures and images for "Sexual dimorphism in the tardigrade *Paramacrobiotus metropolitanus* transcriptome"

### Supplementary Figure 1

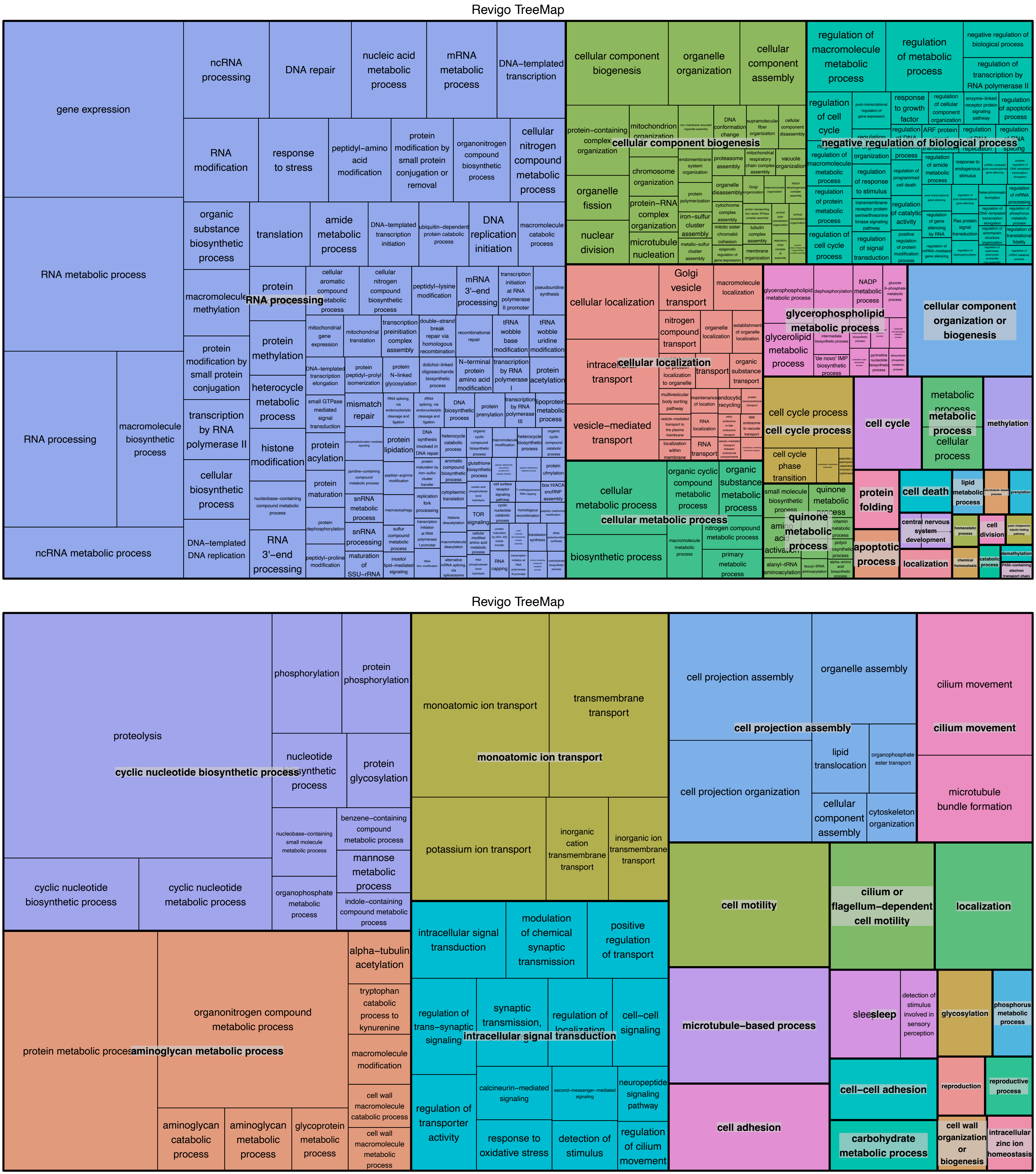

### Supplementary Figure 2

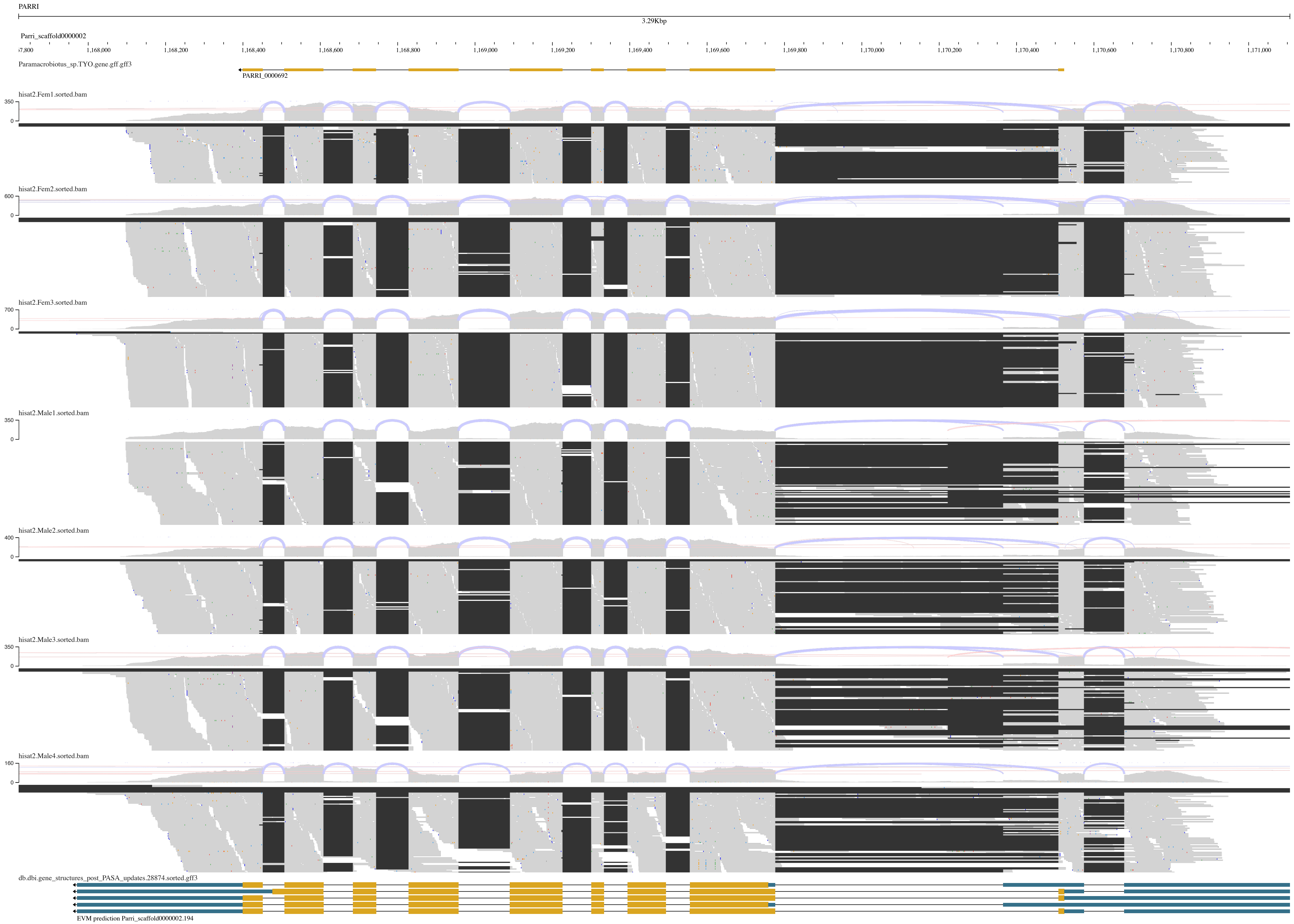

### Supplementary Figure 3

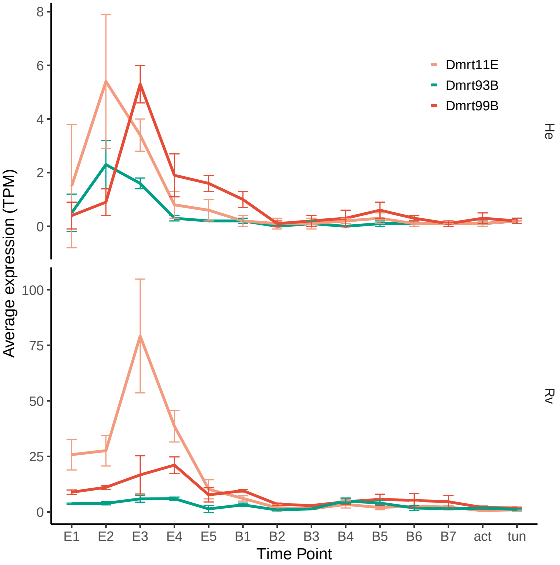

### Supplementary Figure 4

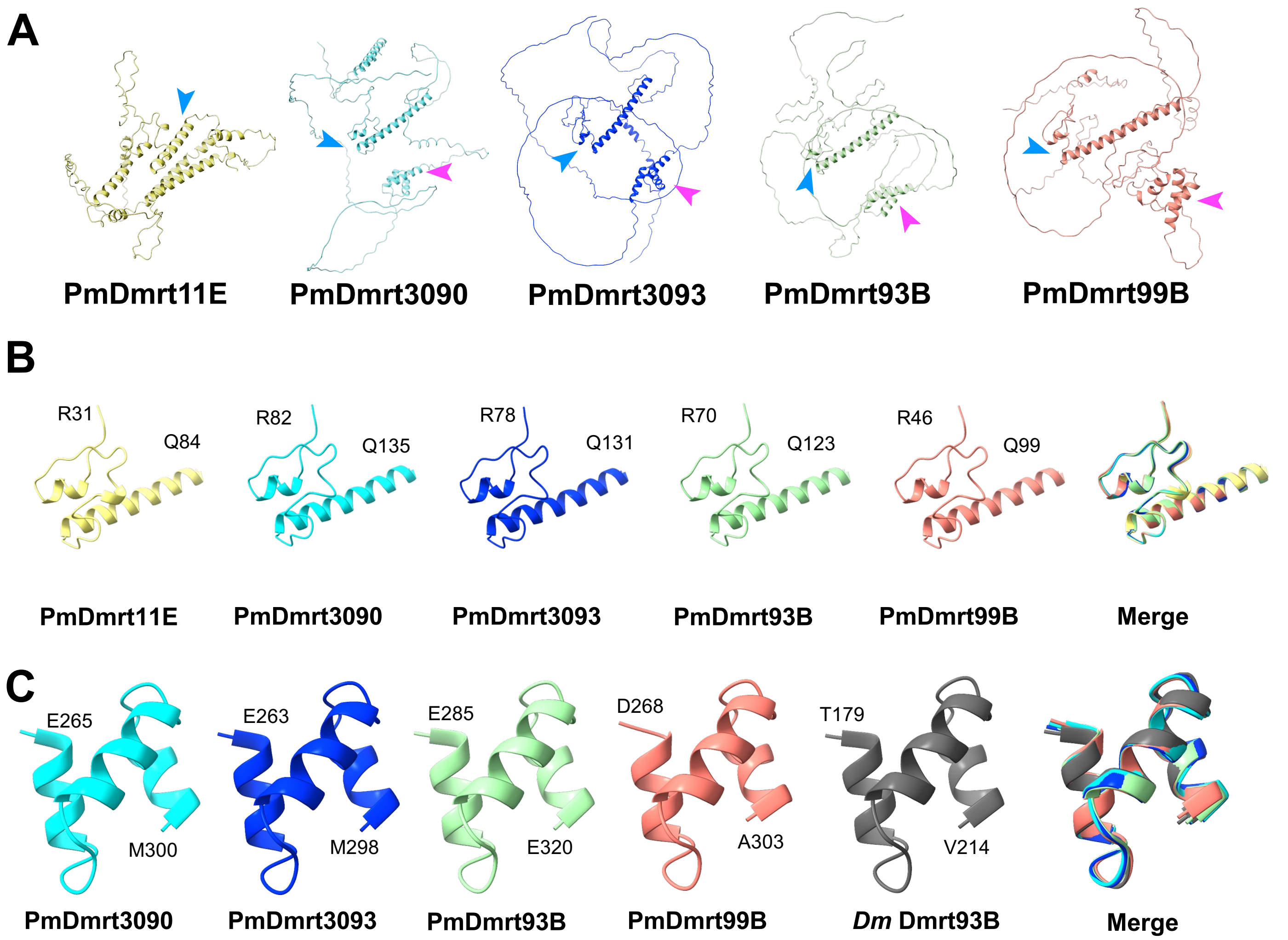
